# Gut microbiome meta-analysis reveals dysbiosis is independent of body mass index in predicting risk of obesity-associated CRC

**DOI:** 10.1101/367466

**Authors:** K. Leigh Greathouse, James Robert White, R. Noah Padgett, Brittany G Perrotta, Gregory D Jenkins, Nicholas Chia, Jun Chen

## Abstract

Obesity is a risk factor for colorectal cancer (CRC), accounting for more than 14% of CRC incidence. Microbial dysbiosis and chronic inflammation are common characteristics in both obesity and CRC. Human and murine studies, together, demonstrate the significant impact of the microbiome on governing energy metabolism and CRC development; yet, little is understood about the contribution of the microbiome to development of obesity-associated CRC as compared to non-obese individuals. In this study, we conducted a meta-analysis using five publicly available stool and tissue-based 16S rRNA and whole genome sequencing (WGS) data sets of CRC microbiome studies. High-resolution analysis was employed for 16S rRNA data using Resphera Insight, which allowed us to achieve species-level information to compare with WGS. Characterization of the confounders between studies, 16S rRNA variable region, and sequencing method, did not reveal any significant effect on alpha diversity in CRC prediction. Both 16S rRNA and WGS were equally variable in their ability to predict CRC. Results from community structure and composition analysis confirmed lower diversity in obese individuals without CRC; however, no universal differences were found in diversity between obese and non-obese individuals with CRC. When examining taxonomic differences, the probability of being classified as CRC did not change significantly in obese individuals for all taxa tested. However, random forest classification was able to distinguish CRC and non-CRC stool when body mass index was added to the model. Overall, microbial dysbiosis was not a significant factor in explaining the higher risk of colon cancer among individuals with obesity.

## Introduction

The percentage of individuals who are overweight or obese in the U.S. has reached epidemic proportions, with the prevalence of individuals who are overweight (32.7%) or obese (34.3%), as defined by body mass index (BMI), in the United States representing about two thirds of adult Americans. The health risks associated with overweight and obesity include diabetes, cardiovascular disease, and cancer. The National Cancer Institute estimates that 3.2% of all new cancers are due to obesity and that 14% of deaths from cancer in men and 20% in women are attributed to obesity (1, 2). Colorectal cancer (CRC) accounts for approximately 142,000 new cancer cases and 50,000 cancer deaths annually, making it the second most lethal cancer in the U.S. (SEER). Several epidemiological studies demonstrate that adult obesity increases the risk of colon cancer 1.2 to 2-fold, with obesity accounting for 14–35% of total colon cancer incidence (1, 3–5). Alarmingly, incidence and mortality from CRC is on the rise among those under the age of 55 (SEER), possibly due to the significant increase in obesity in women (6). For these reasons, it is imperative to identify new methods to reduce the burden of obesity on the risk and mortality from colon cancer. Three areas of inquiry are important for understanding the etiology of CRC: obesity, inflammation, the microbiome.

Several studies indicate that specific microbial taxa are playing a role in the etiology of colon cancer. However, whether the microbiome is also contributing to development of obesity-associated colon cancer in *humans* is completely unknown. One method that has shown promise for identifying early stage colon cancer is through analyzing the microbiome of the gastrointestinal tract (GI). The structure and function of the bacterial community that makes up the human colon, in part, determines the function and health of the colonic epithelium, as well as, the immune system responses. Several studies have found colon cancer-associated microbiota in pre-cancerous colon tissue (adenomas). Further, the microbiome has been used to distinguish pre-cancerous adenomas from CRC, though with variable rates of accuracy (7–9). Several bacteria have been identified as promoters in colon cancer development, including enterotoxigenic *Bacteroides fragilis* and *Fusobacterium nucleatum*(10–14). Both have also been isolated from patients with familial adenomatous polyposis (FAP) or inflammatory bowel disease, which are risk factors for colon cancer (11, 15). Colorectal adenocarcinomas associated with high abundance of fecal *F. nucleatum*, specifically, were found to have the highest number of somatic mutations, suggesting that these mutations create a pathogen-friendly environment (16, 17). In animal models of colon cancer, inoculation of germ-free animals with stool from tumor-bearing animals were found to have more tumors than mice inoculated with stool from tumor-free animals (18). More recently, colonic biofilms from individuals with familial adenomatous polyposis were found to be dominated by *E. coli* and *B*. fragilis biofilms and enriched with genotoxic colibactin and *B. fragilis* toxin (ETBF) genes (11).

Multiple lines of evidence demonstrate that both diet and obesity can significantly alter the microbiome (19–25). One of the first seminal studies illustrating the impact of the microbiome on obesity, transferred the fecal microbiota from monozygotic twins who were obese or lean to germ-free mice. From this study, they were able to recapitulate the obesity phenotype in humanized mice (26). When examining microbiota and subsequent changes in metabolism after fecal transfer from obese mice to germ-free mice, it was found that this obesogenic microbial community had an increased production of SCFAs, which was later shown to abrogate lipid storage (23, 26). Multiple follow-up studies in obese and lean individuals have linked the specific shift in the microbiota to the ratio of *Bacteroides:Firmicutes* (25, 27, 28). However, a recent meta-analysis of these studies indicate that this ratio is not sufficient to differentiate obese from lean individuals in separate studies. Thus, more research is necessary to identify the microbiome-host relationship in individuals with obesity (29, 30).

Chronic inflammation is a hallmark of both obesity and CRC etiology. Obesity is characterized by pro-inflammatory adipose tissue macrophages that secrete high levels of IL-17, a cytokine which is also induced by ETBF in murine models of colon cancer (10, 31, 32). Given the reciprocal relationship between the microbiome and the immune system, it is logical to hypothesize that obesity-associated microbial dysbiosis, combined with a state of chronic inflammation, contributes to the increased risk of colon cancer among obese individuals. In support of this hypothesis, animal models of colon cancer (Apc^1638N^), have demonstrated that a high fat diet or genetically (ob/ob) induced obesity can significantly alter the microbiome leading to a loss of *Parabacteroides distasonis* and an increase in pro-inflammatory factors (22). In a separate model of colon cancer (K-ras^G12Dint^), fecal transfer from high-fat fed mice with intestinal tumors to genetically susceptible mice on a standard diet replicated the disease phenotype (33). Thus, it appears that a high fat diet may be sufficient to change the microbiome into a tumor-promoting community independent of obesity and glucose response. Intriguingly, *Akkermansia muciniphila*, which is reduced in obese individuals and is associated with epithelial barrier function, is paradoxically higher in CRC (34, 35). This data, together with evidence that *A. muciniphila* can modulate glucose metabolism and inflammation in the colon, suggests it may play a role in obesity-associated CRC (35, 36). As these data demonstrate, there are a variety of dysbiotic states that exist in obese individuals, which could further enhance the inflammatory state of the GI tract leading to an increased risk of CRC. No human studies to date have addressed the obesity-associated differences in the microbiome and its relationship to CRC however.

In this study, we utilized multiple publicly available data sets in which either stool or tissue microbiome sequencing was conducted, and from which body mass index (BMI) was also available. Using the bioinformatics tools QIIME (16S rRNA) and Pathoscope (WGS), we processed the 16S rRNA and WGS reads, and derived a taxonomic profile from each of the samples. Furthermore, we inferred taxonomic function to assess potential metabolic differences in obese individuals with CRC. We used these taxa and the metabolic pathway information to determine if a taxonomic signature or if specific taxa were associated with both obesity and CRC. From this analysis, we observed that the dysbiosis associated with obesity was independent from the dysbiosis associated with CRC.

## METHODS

### Sample Population

For this study we identify studies relevant to assess the relationship between obesity and CRC using the microbiome as the independent variable using the following search terms in PubMed (((((“humans”[MeSH Terms] AND (“2006”[PDAT]: “2016”[PDAT])) NOT Review[Publication Type]) AND (obesity[Text Word] OR bmi[Text Word] OR body mass index[Text Word] OR BMI[Text Word] OR obesity[Text Word])) AND (bacterial[All Fields] AND (“microbiota”[MeSH Terms] OR “microbiota”[All Fields] OR “microbiome”[All Fields]))) AND ((“colonic neoplasms”[MeSH Terms] OR (“colonic”[All Fields] AND “neoplasms”[All Fields]) OR “colonic neoplasms”[All Fields] OR (“colon”[All Fields] AND “cancer”[All Fields]) OR “colon cancer”[All Fields]) OR (“colorectal neoplasms”[MeSH Terms] OR (“colorectal”[All Fields] AND “neoplasms”[All Fields]) OR “colorectal neoplasms”[All Fields] OR (“colorectal”[All Fields] AND “cancer”[All Fields]) OR “colorectal cancer”[All Fields]) OR CRC[All Fields]). From our PubMed search we identified 5 (out of 124) studies that met all of our criteria: primary research in a human population, colon or colorectal cancer AND normal stool or tissue collected, raw sequences available from either 16S rRNA or WGS sequencing, body mass index available as a variable in the metadata including age and sex. Together, 5 studies were identified that assessed both BMI and the microbiome in stool or tissue from individuals with adenomas, carcinomas or individuals without disease (Table 1). Three of these studies conducted 16S rRNA sequencing on stool or tissue, and 3 conducted WGS on stool or tissue, with one utilizing RNA sequencing. One study conducted both 16S rRNA and WGS on tissue and stool.

**Table 1:** Summary of obesity, demographic, sequencing, and reads for included data sources.

| Database (Study) | Sample Type | Sequencing Method | Primers * | Species/OTUs<sup>#</sup> | Sample Source (N) | Sample Size by Group |  | Average BMI |  |  |
| --- | --- | --- | --- | --- | --- | --- | --- | --- | --- | --- |
|  |  |  |  |  |  | Obese | Non-Obese | Obese | Non-Obese | P (obese v non-obese)<sup>^</sup> |
| Baxter et al. (2016) | Stool | 16S rRNA | V4 | 9997 | carcinoma(318); control (172) | 118 | 368 | 34.05 | 24.93 | < .001 |
| Feng et al. (2015) | Stool | WGS | NA | 408772 | adenoma (42); carcinoma (41); control (55) | 43 | 113 | 31.98 | 25.62 | < .001 |
| Vogtmann et al. (2016) | Stool | WGS | NA | 356748 | carcinoma (52); control (52) | 13 | 69 | 32.57 | 23.58 | < .001 |
| Zackular et al. (2014) | Stool | 16S rRNA | V4 | 24990 | adenoma (30); carcinoma (30); control (30) | 26 | 56 | 33.84 | 25.09 | 0.002 |
| Zeller et al. (2014)a | Stool | 16S rRNA | V4 | 9969 | control (75); CRC (41) | 19 | 108 | 32.53 | 24.13 | < .001 |
| Zeller et al. (2014)b | Tissue | 16S rRNA | V4/V3-4 | 9988 | carcinoma (48); carcinoma-adjacent (48) | 14 | 80 | 33.3 | 24.5 | < .001 |
| Zeller et al. (2014)c | Stool | WGS | NA | 327491 | adenoma (42); carcinoma (53); control (297) | 34 | 160 | 32.15 | 24.18 | < .001 |

\*1 Primers are NA for WGS

### Processing of Microbial Reads and Calculation of Diversity

All sequence data were downloaded from the NCBI Sequence Read Archive. In order to eliminate differences between studies, we processed the reads using the same methods, either QIIME plus the algorithm Resphera Insight for 16S rRNA sequencing or Pathoscope (v1.0) for processing WGS or RNA-seq reads. For the studies sequencing the 16S rRNA gene, the V4 region was used for all stool samples, as well as, tissue, with the exception of the subsample of tissue from another study used as part of the Zeller et al. 2014 data set. Details regarding sequencing methods and variable regions amplified for each data set are listed in Table 1.

Raw paired-end reads reflecting 16S rRNA fragments were merged into consensus sequences using FLASH (min overlap: 20 bp overlap; 5% max mismatch density), and trimmed for quality (target error rate < 1%) using Trimmomatic and QIIME. PhiX control sequences were identified using BLASTN and filtered. Resulting sequences were evaluated for chimeras with UCLUST (*de novo* mode) and screened for human DNA using Bowtie2 against NCBI *Homo sapiens* Annotation Release 106. Reads assigned to chloroplast or mitochondrial contaminants by the RDP classifier with a minimum confidence of 50% were also removed. High-quality 16S rRNA sequences were assigned to a high-resolution taxonomic lineage using Resphera Insight (37–39) Raw paired-end shotgun metagenomics sequence datasets were also trimmed for quality using Trimmomatic (min final length 75bp) and screened for human genomic DNA using Bowtie2 (--sensitive setting against GRCh38 reference with alternate chromosomes). High-quality passing sequences were submitted to Pathoscope v1.0 for species level characterization (40, 41).

### Prediction of Metagenomic Pathways

We utilized two methods in order to derive abundance of metabolic pathways from the 16S rRNA or WGS sequences. For the 16S rRNA reads, after obtaining the OTU tables, we utilized the PICRUSt algorithm. This method obtains the representative genomes according the nearest neighbor match, and then normalizes the genome abundance using the 16S rRNA copy number for that genome. Once the metagenomics content is binned, it is expressed in terms of KEGG representative ortholog (KO) counts. For the WGS reads, we utilized the HUMAnN algorithm. This method takes as input short DNA or RNA reads and uses BLAST to identify orthologous gene families, which are used to identify metabolic pathways. Once identified, the pathways are then normalized by presence/absence of the taxa, and additionally by relative abundance of the taxa present in the sample. These data were used for downstream statistical analysis to compare obese and normal stool samples from individuals with or without CRC.

### Statistical Analyses

Prior to analysis we rarefied the data to the sample with the lowest number of reads. In order to test the association between BMI and the microbiome, we grouped our statistical analyses into four subgroups: A) normal stool samples (healthy controls), B) CRC stool or CRC tissue, and C) pooled samples (healthy controls and CRC), all of which were adjusted for age and sex. Group C was further adjusted for disease status.

For alpha diversity measurements, we used both the observed number of OTUs and the Shannon Index. To determine associations with BMI, we treated it as a continuous variable (as a covariate) in the main analysis. For additional analyses, we also dichotomized the subjects into non-obese (BMI < 30) and obese (BMI >= 30) according the WHO guidelines.

For beta diversity measurements, we utilized four distance measurements unweighted UniFrac, weighted UniFrac, generalized UniFrac and Bray-Curtis for 16S datasets. For WGS/RNA-seq data, where we do not have the phylogenetic tree, we instead used two non-tree-based distance measurements Jensen-Shannon and Bray-Curtis (42). Different distance measurements represent different views of the microbial community and multiple distance measurements are used to have a more comprehensive view. In order to determine the difference in community membership between BMI categories, we used the PERMANOVA test on single distance measures, with the omnibus test on the combination of all distance metrics (PermanovaG, ‘GUniFrac’ R package) (43).

In order to compare taxonomic abundance between groups, we used as input OTU counts. Negative binomial regression was used with BMI as a continuous variable for analysis of the microbiome while controlling for age and sex. Using multilevel modeling, the effects of confounders in study designs are examined. In this multilevel model, the study is defined as level 2 and the individual observations are level 1. At level 1, the outcome is CRC status (1=has CRC, 0=does not) and is predicted by an intercept, alpha diversity and BMI. At level 2, the level 1 regression coefficients (i.e. β_0j_, β_1j_, and β_2j_ for the intercept, alpha diversity and BMI regression coefficient, respectively) are modeled by the study characteristics. In this model, the level 1 regression coefficients vary among studies, which means, for example, that the effect of alpha diversity to predict CRC status varies by study and study characteristics. For precisely, we are estimating the following model:

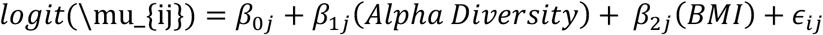

Where the regression coefficients are modeled by study characteristics. For example,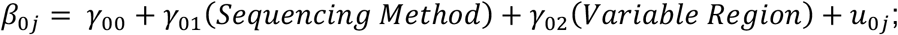which defines how the model parameter vary by study characteristics. In this model, the *γ*’s represent the level two model parameters and u is the study specific error term. Estimation of this model is employed using the lme4 (linear mixed effects) package in R (44). Due to the few number of studies included in this meta-analysis, the estimation of the variance of the level 1 parameters is uncertain and should be interpreted with caution.

In order to compare taxonomic abundance between groups, we used as input OTU counts. Negative binomial regression was used with BMI as a continuous variable (obese vs. non-obese) for analysis of the microbiome while controlling for age and sex. Multiple testing procedure was conducted on these values using BH-based false discovery rate control. The criterion to declare significance was q-value <0.2. Furthermore, the abundance of *Bifidobacterium catenulatum* was examined among groups of obese or non-obese individuals with and without CRC. The standardized mean differences among studies was calculated using Hedge’s g, a bias corrected estimate of standardized mean differences. Estimation was employed using the meta package in R (45).

Mediation analyses were also conducted. The goal of these analyses is to uncover if the relationship between BMI and CRC status is mediated by bacteria present. First, bacteria were dummy coded for presence or not for everyone. By dummy coded solely for whether an individual has a given bacteria or not, these results are not meant to show mediation among varying levels of each bacterium. Second, the relationship between BMI and CRC status was estimated by using a simple logistic regression model. Third, the classic mediation model was estimated by using the lavaan package in R (46). This model is estimated for the presence of each bacterium. Lastly, the change in the odd ratio is calculated between models. The change in the OR is an estimate of the mediation effect that a bacterium has on the relationship between BMI and CRC status.

Further exploration of whether taxonomic abundance among obese or non-obese individuals is indicative of CRC utilized random forest analyses. Random forest analysis is a machine learning/predictive modeling algorithm designed to estimate an ensemble of decision trees that are combined to give an estimate of an output. In this study, we employed random forest analyses as a classification of obesity (obese vs. non-obese) conditional on the status of CRC. Four random forests were grown for each study dataset when possible; the forests were grown using the relative abundance of taxa with or without age and sex included at the OTU and genus level for two subsets of data that were conditioned on CRC status (CRC or adenoma). The resulting models aimed to classify individuals as obese based on the microbiota composition, and these classification models were tested with 10-fold cross validation. The receiver-operating-curve (ROC) of these classifications was also inspected for how sensitive the models are to detect obese individuals and how specific these models are to select only individuals that are obese. A measure of model quality is the area under curve (AUC), or area under the ROC, where an AUC of one is perfect prediction and an AUC of .5 is pure chance or prediction. Another benefit of using random forest analyses is that an estimate of the predictive importance of each OTU or genera is estimated. This estimate of importance is found by the predictive quality of model conditional of the ensemble trees that do not contain that specific input variable (OTU or genera in this case). All processed data and code for this analysis has been deposited at:https://github.com/GreathouseLab/CRC_BMI_meta_analysis.

## Results

### Database and study selection

We performed a systematic review and meta-analysis guided search of the literature. Within this search we included studies that analyzed the microbiome of the stool or tissue from patients with colon cancer, and which also had clinical information from patients on BMI. From this initial search, we identified 24 studies. After eliminating studies in which BMI information could not be obtained, 5 studies were included in the final analysis (Table 1). Given that our central hypothesis is predicated on a difference in microbial structure and composition between obese and non-obese individuals, we focused our initial analyses on the Baxter et al. study, which has adequate sample size to detect differences between these two groups (8). The remaining studies were used as comparators to support or negate any associations found between the microbiome and obesity.

### Characterization of cofounders between studies

A major issue facing microbiome studies is the lack of standardized methods for collection, storage, nucleotide extraction, sequencing methodology and bioinformatic analysis. Thus, we began our analysis by characterizing the effect of 16S rRNA variable region and sequencing methods (16S rRNA or WGS) on observed OTUs and Shannon diversity on prediction of CRC. Unfortunately, we could not fully test the effect of nucleotide extraction as the Feng et al. study did not provide this information. We chose to focus on alpha diversity for this analysis given that it is a low-resolution measure, which allows for comparison across studies. Using multilevel modeling to predict CRC status we calculated the average log2 OR (logit) for each study when these level 2 predictors (variable region and sequencing method) are included in the model. The results of this analysis demonstrated that alpha diversity and obesity vary by study but do not significantly change the probability of having CRC (Figure 1A-B; Figure S1). Interestingly, the Feng et al. data set display an unusual inverse relationship between CRC ad BMI that strongly impacts prediction of CRC, possibly due to geographic and dietary differences in this population. Since all but one of the studies used the V4 16S rRNA region, it was difficult to determine if this variable had a significant impact. Between the studies that used different extraction techniques, Zeller (GNOME DNA Isolation Kit, MP Biomedical) vs Baxter and Zackular et al.(Power Soil, Mo Bio), we did not observe an effect of extraction technique on the relationship between alpha diversity and probability of CRC (Figure 1A-B). Further, the predictive ability of 16S rRNA data, alpha diversity, to classify CRC varies among studies but using WGS does not improve this predicative ability nor does variable region choice (Figure 1A-B). Overall, among the potential confounders we tested, we did not observe a significant effect on the ability of alpha diversity to classify CRC cases and controls when controlling for obesity.

**Figure 1:**
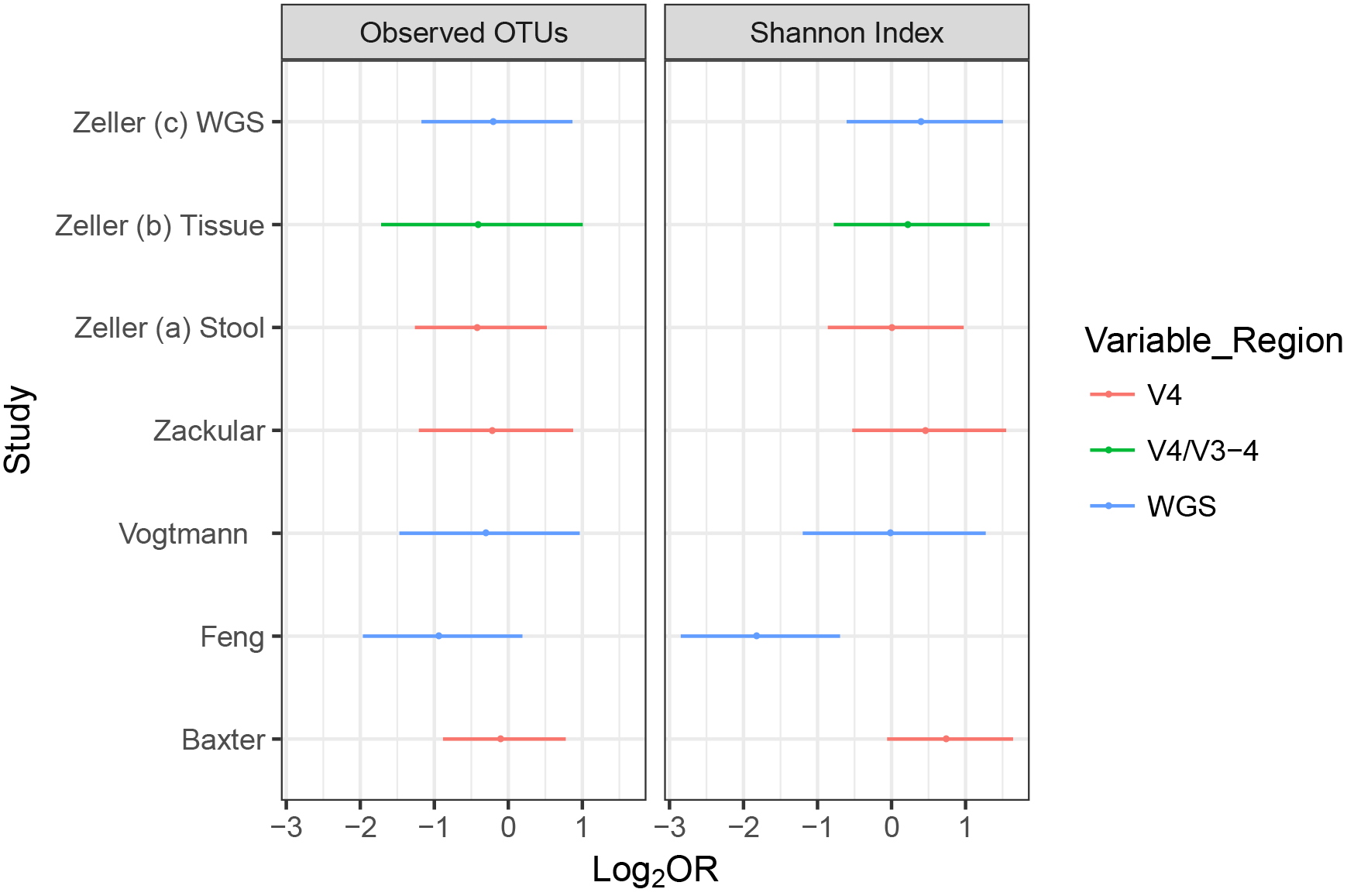
Variance in ability of alpha diversity to predict odds (log2) of CRC controlling for obesity and study confounders. The Log_2_ odds ratio of CRC using observed OTUs (left panel) or Shannon Index (right panel) as predictors. The multilevel model includes obesity (level 1), and sequencing method (16S rRNA or WGS) and variable region (V4 or V3–4) (level 2) as coefficients.

### Alpha Diversity Analysis

We next sought to validate previous studies showing differences in alpha diversity between obese and non-obese individuals without CRC. In order to analyze alpha diversity within each sample study population, we calculated both richness, observed OTUs, and Shannon diversity, which considers both evenness and richness. We conducted linear modeling analysis using BMI as a continuous measurement and calculated the observed OTUs and Shannon diversity controlling for age and sex. Confirming previous microbial studies of stool from healthy (non-CRC) individuals (30), we also found significantly lower Shannon diversity in individuals that are obese without cancer from two of the 16S rRNA data sets (Baxter and Zeller et al. (WGS)) and lower richness in the Zeller et al. (16S) data; unadjusted Mann-Whitney U tests did not show this same result comparing individuals with and without obesity (Fig. 2A; Supplemental Fig S2A and Table 2). Supporting previous meta-analyses, however, studies with N<100 subjects displayed similar trends but did not reach statistical significance. When we asked if this same trend of lower Shannon diversity was present in obese individuals with CRC, we saw no association, with the exception of the Feng dataset, which demonstrated a significantly higher alpha diversity with higher BMI both as continuous and categorical models, but not in the unadjusted analysis (Fig. 2B; Supplemental Fig S2B and Table 2). These results may be due to geography and diet of Asian populations. We chose not to analyze the *Bacteroides/Firmicutes* ratio as this has not demonstrated to be a consistent measurement of predicting obesity in human studies (30). Together, these data indicate that while there is an association between community composition and obesity in those without CRC, this association is not present in those with both obesity and CRC.

**Table 2:**
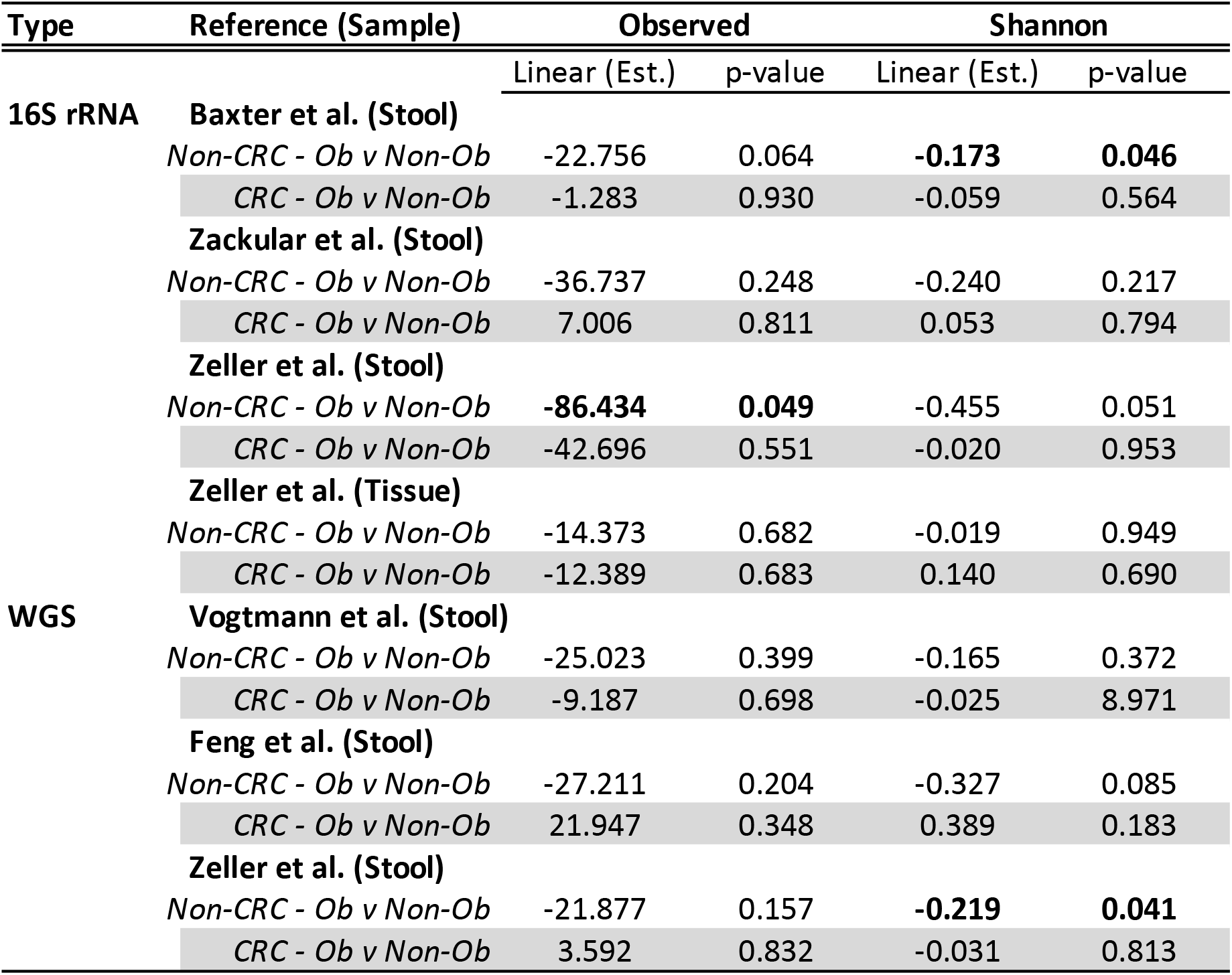
Summary of Alpha-Diversity Analysis.

**Figure 2:**
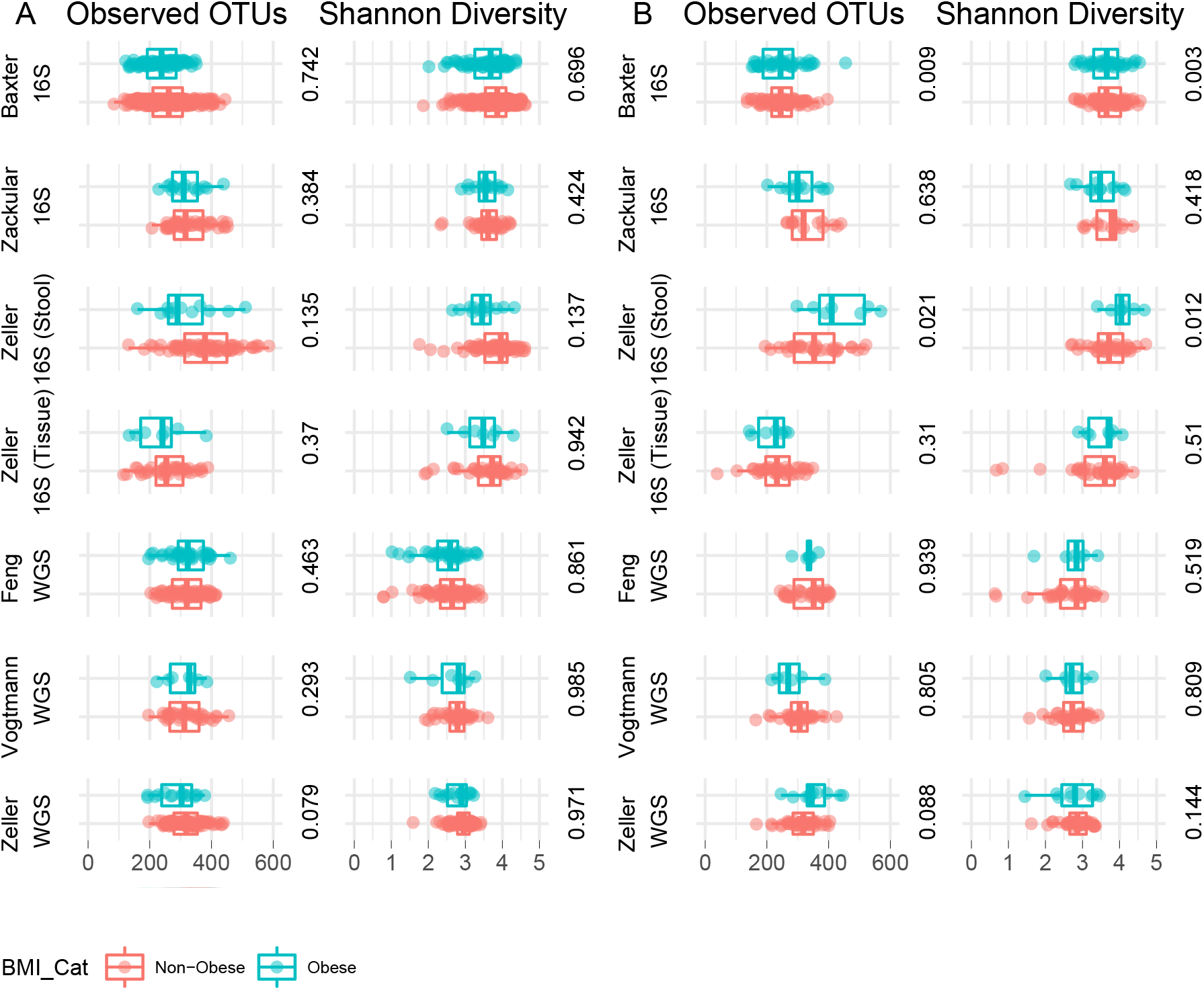
Alpha diversity in individuals with or without obesity and with or without CRC A) Observed OTUs and Shannon diversity in individuals without CRC B) or with CRC comparing individuals with or without obesity. Reporting p-values are from Mann-Whitney U Test comparing the alpha diversity of individuals with or without obesity.

### Beta Diversity Analysis

We next asked whether we could detect microbial community differences in structure between obese and normal weight individuals with or without CRC. In order to conduct this analysis, we calculated the distance matrix for each study using UniFrac or Bray-Curtis (BC) for 16S rRNA datasets, and BC or Jaccard-Sorrensen (JS) distance for WGS datasets. Further, we calculated the omnibus p-value for comparison of all distance matrices (47). In all of the data sets analyzed, except Vogtmann et al. (WGS), Zeller et al. (WGS), and Zeller et al. (16S rRNA/tissue), we observed a significant difference (omnibus p-value <0.05) in community structure between obese and non-obese individuals without CRC (Table 3; see Fig. S3A in supplemental materials). This same analysis in individuals with CRC (obese v non-obese), however, yielded only one significant observation in the Feng et al. dataset (Table 3; Supplemental Fig. S3B), supporting the observations with community composition. Thus, similar to community composition, community structure is associated with BMI in individuals without CRC but not in those with CRC.

**Table 3:**
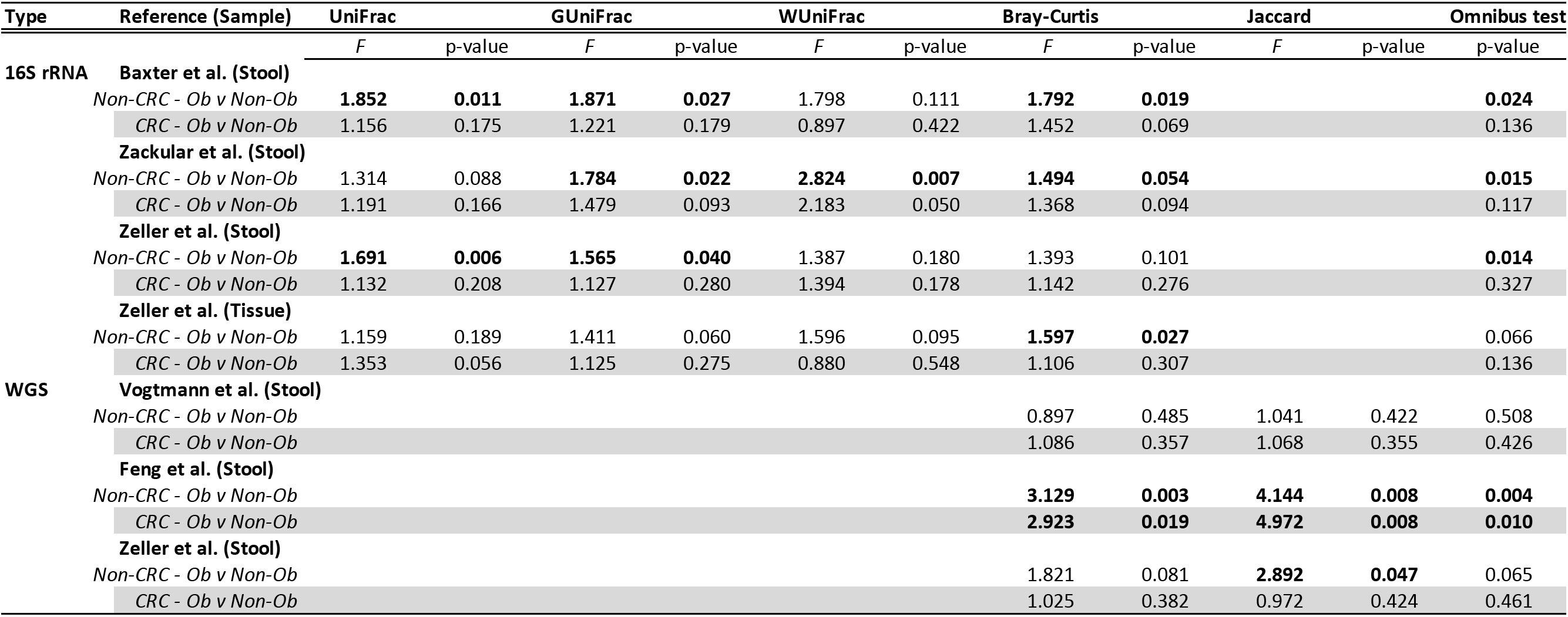
Summary of Beta-Diversity Analysis.

### Taxonomic Diversity Analysis

Again, we began our taxonomic analysis comparing individuals with and without obesity among individuals without CRC as a means of validating previous studies, using as our reference the largest study dataset, Baxter et al. (8). From this analysis, controlling for age and sex, a significantly lower relative abundance of several *Ruminiococcus spp*. was identified in the two of the datasets (Zackular et al., Zeller et al. (16S rRNA stool), Zeller et al. (WGS), as well as, *Coprococcus spp*. (Baxter et al., Zackular et al., Zeller et al. (16S rRNA stool)), *Bacteroides spp*. (Baxter et al., Zackular et al., Feng et al., Vogtmannn et al., Zeller et al. (WGS)), *Bifidobacterium spp*.(Zeller et al. (WGS)) and *Akkermansia muciniphila* (Zackular et al., Zeller et al. (WGS)) (Supplemental Fig. S4 and Supplemental Table 1). When combining all differentially significant species, those from genus *Bacteroides* and *Bifidobacteria* appeared most often to differentiate individuals with and without obesity (Supplemental Table 1). While no one genera or species was found to be differentially abundant (higher or lower) between all 5 datasets comparing individuals with or without obesity among individuals without CRC, the genus *Bacteroides* contained the greatest number of differentially abundant species in individuals with obesity in all but one dataset (Supplemental Table 1).

### Mediation effect of differentially abundant taxa on obesity-associated CRC classification

In order to determine if any taxa were affecting (mediating) the relationship between BMI and CRC probability, we took two approaches. The first approach was a classical mediation test, in which we constructed three tests. First, we estimated the odds-ratio (OR) of individuals with higher BMI being more likely to be classified as having CRC. Second, we estimated the same relationship between BMI and CRC status while controlling for the mediating effect of differentially abundant bacteria. Meaning, if the bacterium mediates the relationship between BMI and CRC probability then the OR for BMI will decrease. Third, we calculated how much *change* in the OR occurred from the first to second model. Thus, from this change in ORs, we estimated how much of an effect including each taxa had on increasing or decreasing the probability of being classified as CRC for each one unit increase in BMI. From this analysis, we identified several taxa that increased or decreased the probability of CRC (Supplemental Tables 2–3). Species from the *Bacteriodes, Ruminococcus* and *Prevotella* genera, as well as, *Bifidobacterium catenulatum* decreased the probability of CRC with increasing BMI, except for two species of *Prevotella* which increased CRC probability. The mediation effect of these taxa, however, was relatively weak; less than 1% change in OR (change in probability of CRC, OR range = –9e-05 – –0.01) (Supplemental Table 2), with the majority showing a negative effect and only 8/34 showing a positive effect; none showed a significant mediation effect (Supplemental Table 2).

In our second approach, we derived an overall mediation effect using the FDR adjusted p-values (q-values) from our analysis of the Pooled BMI data (association between BMI and microbiome using all samples only, adjusting for disease status, sex and age) and Pooled DS data (association of disease status with microbiome using CRC and normal samples, adjusting for BMI, sex and age). These overall Q-values were approximately based on 1 – (1 – q1)*(1-q2), where q1 and q2 are q values for the BMI and DS associations on the pooled data set (q value can be interpreted as the probability of being false positive, 1 – (1 – q1)*(1-q2) is the probability of being false positive in either of the associations, assuming independence between the two tests). The q-values were calculated for each data set, and q-values for taxa <20% were considered to have a significant mediating effect. Using this approach, we looked for taxa that had a significant mediating effect between studies and identified two, *Phascolarctobacterium succinatutens* and *Streptococcus salivarius*; however, they were only shared between 2/6 studies each (Supplemental Table 4). Overall, these results indicate the majority of bacteria associated with CRC and BMI decrease the odds of CRC in individuals with obesity, but only weakly.

In addition, to determine if previously identified CRC-associated taxa, *F. nucleatum, F. prausnitzii, B. fragilis*, or *A. muciniphila*, were altered in individuals with obesity in their ability to differentiate CRC from non-CRC, we calculated the log2 odds ratios for each species. (Supplementary Fig. S5). Overall, among individuals with obesity, *F. nucleatum* consistently showed stronger prediction (log2 OR) of CRC.

### Ability of the microbiome to classify obesity-associated CRC

Given that previous studies have demonstrated the predicative capability of the microbiome in generating classifiers for CRC, we next asked whether a taxonomic consortium could accurately predict obesity-associated CRC. Using the machine learning method random forest, we calculated importance scores among obese individuals at the OTU or genus level using 10-fold cross-validation in individuals with adenomas or CRC. These values were then used to calculate area under the receiver operator curve using age and sex as co-variates or the microbiome alone. Among all obese individuals, the average of all AUC values predicting CRC cases at the OTU and genus level was 0.66 (0.47–0.84) and 0.68 (0.47–0.94), respectively (Fig. 3B). Similarly, among obese adenoma cases, average AUC values at the OTU and genus level were 0.61 (0.48–0.86) and 0.60 (0.52–0.73), respectively (Fig. 3A); demonstrating high heterogeneity among studies in predicting CRC or adenomas in obese individuals. Lastly, we sought to validate CRC classifiers developed by Baxter et al. and Zeller et al. by agnostic application of our random forest classifier on each dataset using all genera or OTUs. While Zeller et al. used a more complex statistical approach to construct their classifier, we choose to apply the same method (48) to each study for the purposes of comparison, which was almost identical to Baxter et al. (excluding smoking and hemoglobin test results). Overall, the microbiome by itself or controlling for BMI, age and sex, had low and variable AUC values (OTU; AUC=0.53–0.79; Genus; AUC=0.59–0.81) in most studies. We were able, however, to validate the classifier from the Baxter et al. and Feng et al. studies; our AUC values were 0.79 (Baxter et al.) and 0.81 as compared to Baxter et al. (AUC=0.84) and Feng et al. (AUC=0.96). Although we could not approach the classifier values from the Zeller et al. study (AUC=0.84; without FOBT), this was likely due to the difference in their approach in building the classifier. In general, these data indicate that the microbiome together with clinical data, and likely FOBT or similar tests, could have diagnostic utility.

**Figure 3:**
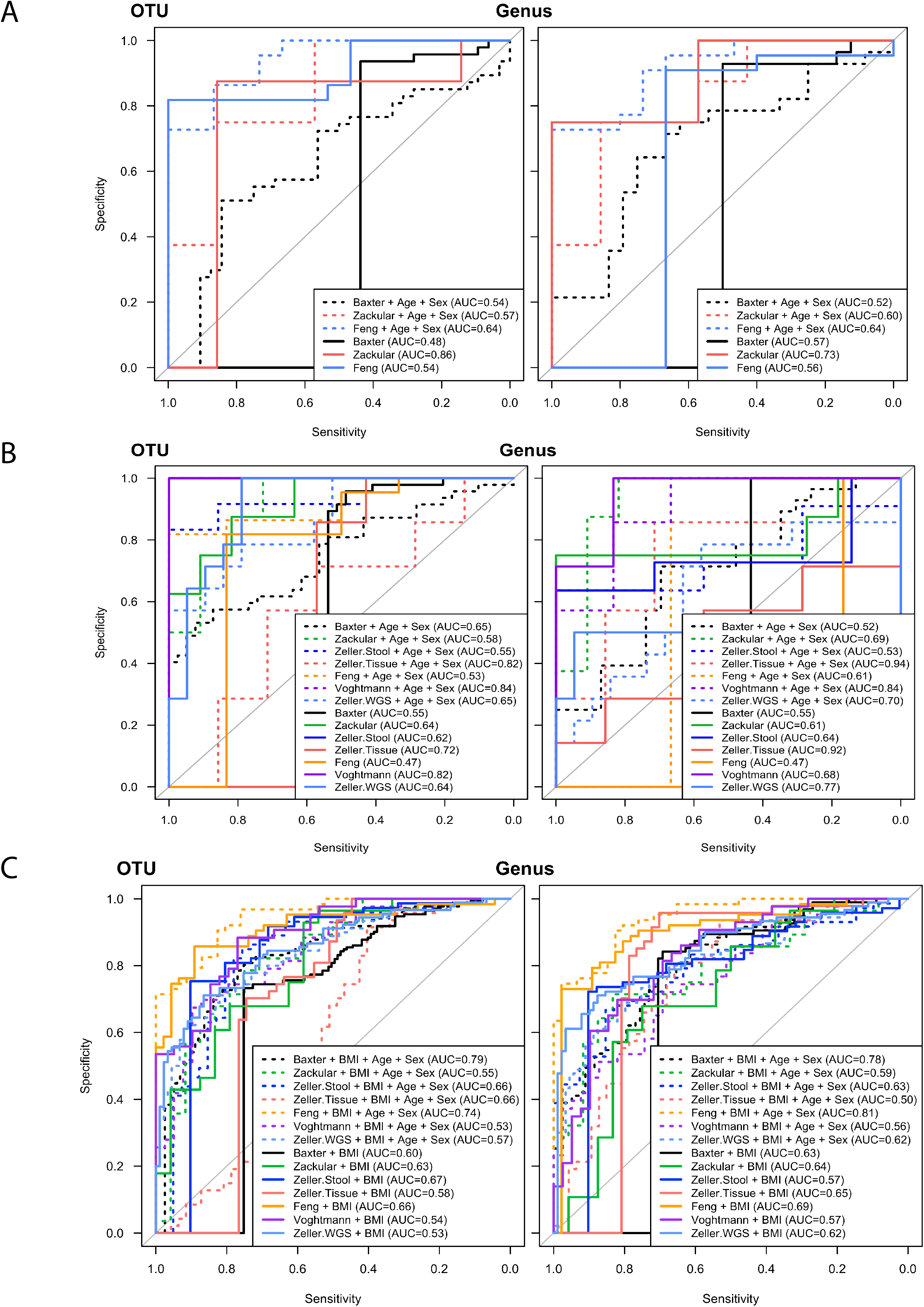
Microbial classifiers of CRC and obesity-associated CRC. A) Receiver Operating Curve (ROC) for the random forest classification analyses for obese vs. non-obese in individuals with CRC for each study. AUC is the 10-fold cross validated area under the curve. B) ROC for the random forest classification analyses of obese vs. non-obese in individuals with adenomas for each study. Due to a lack of cases with adenomas in some studies a random forest was not possible and are therefore not shown. C) ROC for the random forest classification analyses of CRC vs non-CRC in each dataset adjusted for BMI, age and sex.

## Analysis of Inferred Taxonomic Function

Multiple studies have demonstrated that taxonomic abundance alone does not accurately reflect the metabolic function of the entire community. Thus, we interrogated the metabolic potential of the bacterial community using the bioinformatics tool PICRUSt in order to obtain predicted functions. Among individuals without CRC, we identified biotin synthesis and biotin metabolism inversely correlated with BMI in the Vogtmannn (WGS) stool samples, and the urea cycle (M0029) inversely correlated with BMI in both Vogtmannn (WGS) and Feng (WGS) (Figure 4A-B). However, none of these predicted functions differentiated obese individuals among all studies. When we conducted this analysis in individuals with CRC, no shared correlations were identified when comparing obese and non-obese. Predicted functional analysis therefore, did not further distinguish obesity-associated CRC from those with CRC and normal BMI.

**Figure 4:**
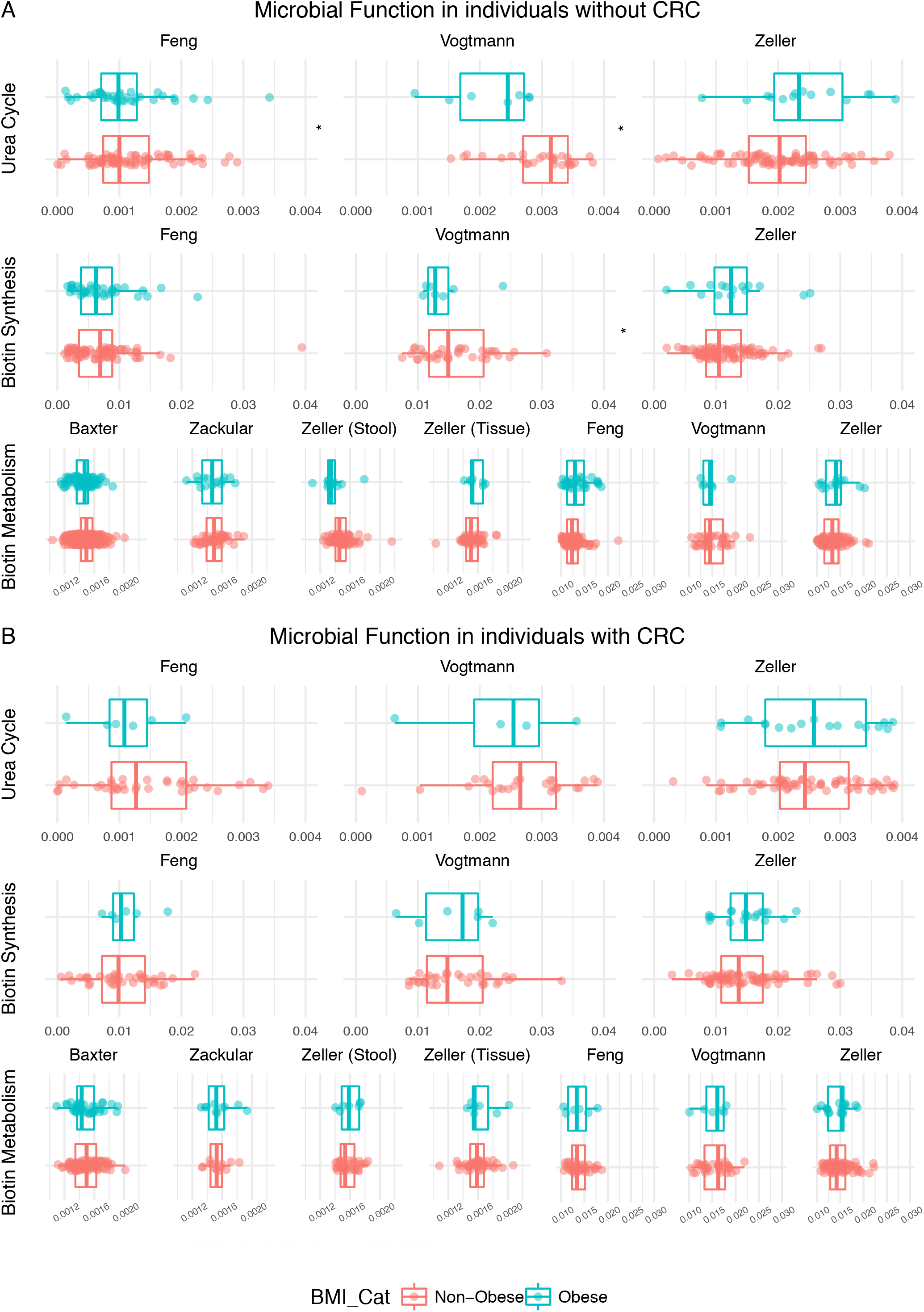
Pathway abundance analysis in individuals with or without obesity among individuals with or without CRC. Relative abundance of KEGG metabolic pathways (16S rRNA) or modules (WGS) inferred from PICRUSt or HUMAnN, respectively. Significance was calculated using the Wilcox test correction for multiple hypothesis testing; asterisks are representative of significance at adjusted p-value <0.2.

## Discussion

Evidence clearly demonstrates an intimate link between inflammation, obesity, and the microbiome (34, 36, 49–56). *In vivo*, multiple studies indicate an interaction or mediating effect of the microbiome in promoting colon tumorigenesis in the presence of a high-fat diet or genetic-induced obesity (21, 22, 57–59). In this study, using BMI as a measure of obesity, we were able to initiate the first analysis addressing this outstanding question in human subjects.

This is the most comprehensive high-resolution study of the microbiome in individuals with and without obesity among those with CRC, using multiple sequencing platforms and methods. In this meta-analysis, we describe both obesity- and CRC-associated results. First, we found both community structure and composition in stool and tissue samples from individuals with CRC are independent of BMI. Second, we identified a weak effect of the majority of species associated with both BMI and CRC on risk of CRC. Lastly, we show the microbiome, by itself or modeled with age and sex, is insufficient to classify adenomas or CRC from obese controls. However, when controlling for clinical variables and BMI, we are able to achieve similar levels of CRC classification to other studies (48). Overall, by combining species-level resolution from 16S rRNA and WGS data, we were able to define the microbial community structure and function at a high resolution, revealing overall a weak effect of the microbiome on mediating CRC risk among individuals with obesity as compared to those with normal BMI.

While this study did not identify any strong universal BMI-associated microbial biomarkers of CRC, many mechanisms are likely key in driving the increased risk of CRC in obese individuals that we could not account for in this study. These include tumor location (left vs right), mutation profile, differentiation, mismatch repair status, and diet; some of which have shown to differentiate individuals with obesity among those with CRC (60–63). A high fat diet may be more important than BMI or obesity in driving the deleterious changes in the microbiome in individuals with obesity. In support, feeding a high-fat diet to K-ras^G12Dint^ mice is sufficient to drive tumorigenesis from 30% to 60% (33). Moreover, when feces from high fat fed mice (K-ras^G12Dint^) are transferred to healthy (K-ras^G12Dint^) mice, tumor burden is increased along with diminished immune cell recruitment (33). This was prevented, however, when supplemented with butyrate, which also increased *Bifidobacterium* abundance as compared to mice not supplemented with butyrate (33). We also found several species of *Bifidobacterium* lower in individuals with obesity among those with and without CRC. Interestingly, butyrate and butyrate producing bacteria were shown to be increased in African-American men after switching to a traditional high fiber, low-fat rural African diet (64). Again, similar to the results of high-fat feeding promoting CRC, which was abrogated with butyrate treatment, the aforementioned study found that the high-fat Western diet of African-Americans was associated with higher secondary bile acids, known promoters of carcinogenesis. Together, these studies indicate that a high fat diet, specifically from saturated fats, may be interacting with the microbiome to create a pro-inflammatory environment conducive to colon carcinogenesis.

Other possible mechanisms explaining the increased risk of CRC in individuals with obesity include a lack of balance in key immune regulatory cells, specifically regulatory T cells (Tregs) and B lymphocytes. Demonstration that Tregs are important in promoting colon tumorigenesis, indicates that species that can control their activation may be important in controlling CRC development. Specifically, when *B. pseudocatenulatum* (CECT 7765) was given orally to obese mice, it increased Tregs and reduced pro-inflammatory cytokines (IL-17A and TNF-a), which further supports the hypothesis that certain species may protect against chronic inflammation and development of CRC (51). In reports measuring dietary inflammatory factors (empirical dietary inflammatory pattern), individuals with higher inflammatory scores had fewer tumor-associated adaptive anti-tumor immune cells suggesting immune evasion (65). Moreover, using this same approach, higher inflammatory scores were associated with higher tumor-associated *F. nucleatum* in CRC (66). Again, these findings support a distinct influence of diet on the microbiome in CRC development apart from obesity.

BMI is crude measure of obesity, and other more accurate measures (e.g. waist circumference, adipokines, etc.) are required to fully explore the relationship between obesity, inflammation, and the microbiome in development of CRC. An exemplar of this relationship is demonstrated for lung cancer, wherein the use of BMI demonstrates a lower risk of lung cancer is associated with higher BMI but use of waist circumference or waist to hip ratio demonstrates and increased risk of lung cancer (67). Thus, this study sets the stage for future research to consider adding measures of adiposity beyond BMI when studying the etiology and risk of CRC, as well as, other cancers influenced by obesity.

This study has several strengths, as well as, limitations. One important strength, was the ability to use multiple peer-reviewed studies that had similar study designs and sequencing methods. As the Microbiome Quality Control Project demonstrated, multiple factors (e.g. DNA extraction method) can contribute to differential findings between studies, and thus our ability to control for these confounding factors reduced this bias (68–71). Also, the ability to confirm the presence of multiple taxa using separate sequencing methods, 16S rRNA and WGS methods, further strengthened the design of this analysis. The limitations of this study include small sample sizes in the majority of studies, use of only one anthropometric measurement of obesity and lack of other informative factors including dietary fat intake, previous weight loss prior to CRC diagnosis, microbial metabolites and biofilm presence. Sample size is a key limitation when looking at the relationship between obesity (BMI) and the microbiome as previously demonstrated (30). Only the Baxter (8) study was sufficiently powered to detect a significant difference in the microbial community between normal and obese individuals. While we were able to identify taxa that differentiated obese and normal individuals with CRC in this study specifically, these taxa were not consistent across all studies, indicating that other factors such as metabolites, biofilm or the immune system are stronger contributors to this relationship. While we were able to derive inferred function from the microbial sequences, without more intensive direct measurement of the metabolites (i.e. mass spectrometry), we cannot fully assess these differences. Additionally, studies have illustrated this lack of relationship between specific taxa and CRC, and instead identified a stronger association with the presence of biofilm formation. Lastly, animal studies demonstrating that CRC promotion by a high fat diet was independent of obesity supports our findings and suggests that dietary fat has a greater impact than obesity on the microbiome and its tumor-promoting capacity in CRC etiology. This will therefore be important to consider in the obesity-CRC relationship in future research.

Overall, our validation of microbiome-based classifiers indicates this approach, in combination with FOBT or FIT tests, is well supported for continued development. More important, these data along with other studies indicate that diet, rather than obesity, is creating a pro-inflammatory microbial community increasing CRC risk. Hence, characterizing the role of the diet in addition to the microbiome in CRC etiology is necessary, which will require more detailed molecular analyses and well-designed longitudinal human studies to identify early stage dietary and microbial biomarkers prior to disease.

## Author contributions

KLG conceived of the study, analysis plan, analyzed data, interpreted results and participated in writing and review; JW downloaded and processed all sequencing data; JC, GDJ and NP conducted statistical analyses; BGP processed data; JW, KLG, JC, GDJ, NP, BGP, and NC provided technical and data interpretation assistance and manuscript review. All authors read and approved the final manuscript.

## Acknowledgements

We would like to thank the participants in each of the studies for their time dedication to supporting colon cancer research. Also, we thank all of the authors from each study for making their data publicly available for analysis, and to allow for transparency and reproducibly. Our thanks to the members of the Baylor Writing Group, Joe Taube, Karen Melton, Elise King, Elyssia Gallagher, for their feedback and editing.

## Supplemental Figures

**Figure S1:**
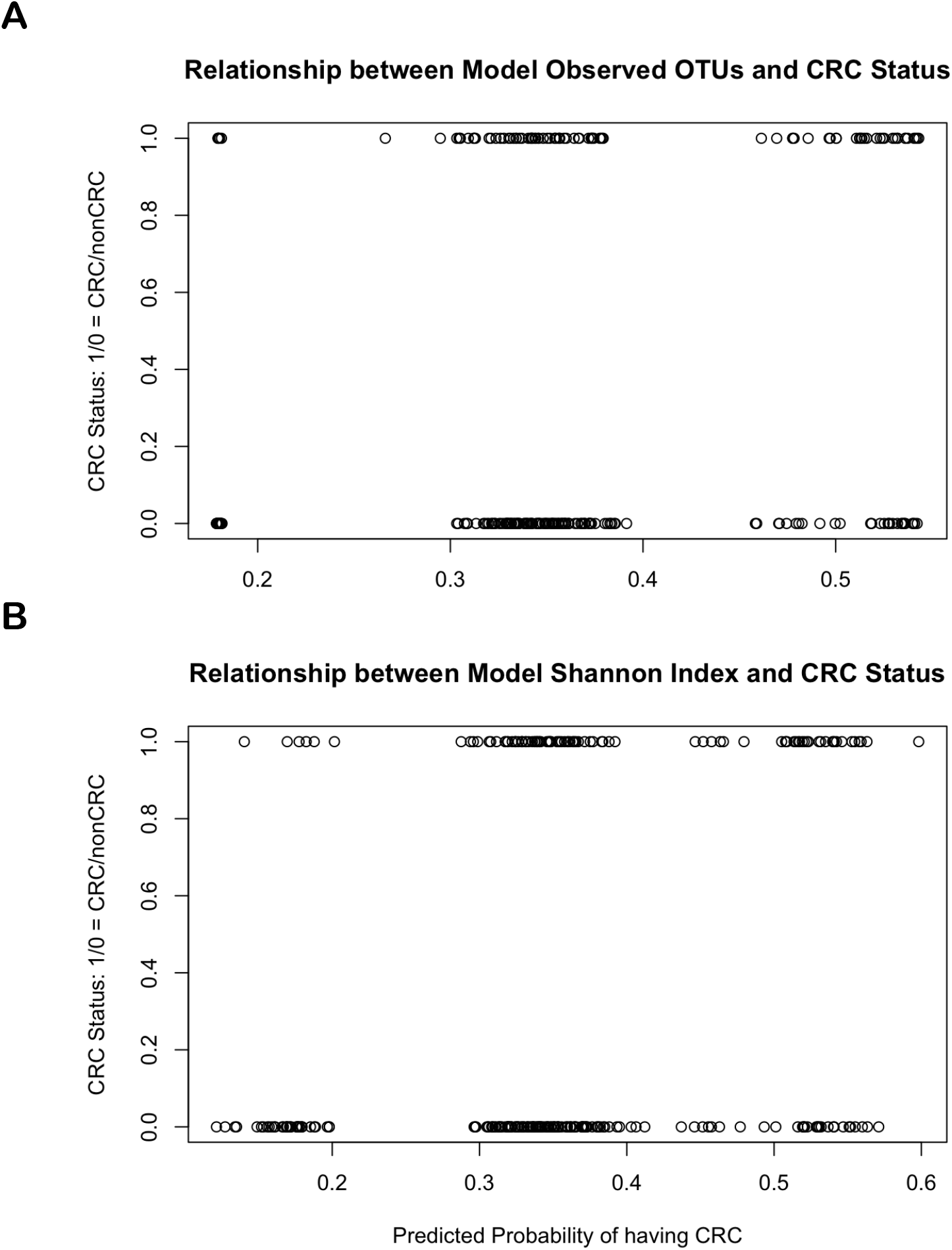
Probability of having CRC using alpha diversity as a predictor among individuals with obesity. Predicted probability of having CRC using A) observed OTUs or B) Shannon Index.

**Figure S2:**
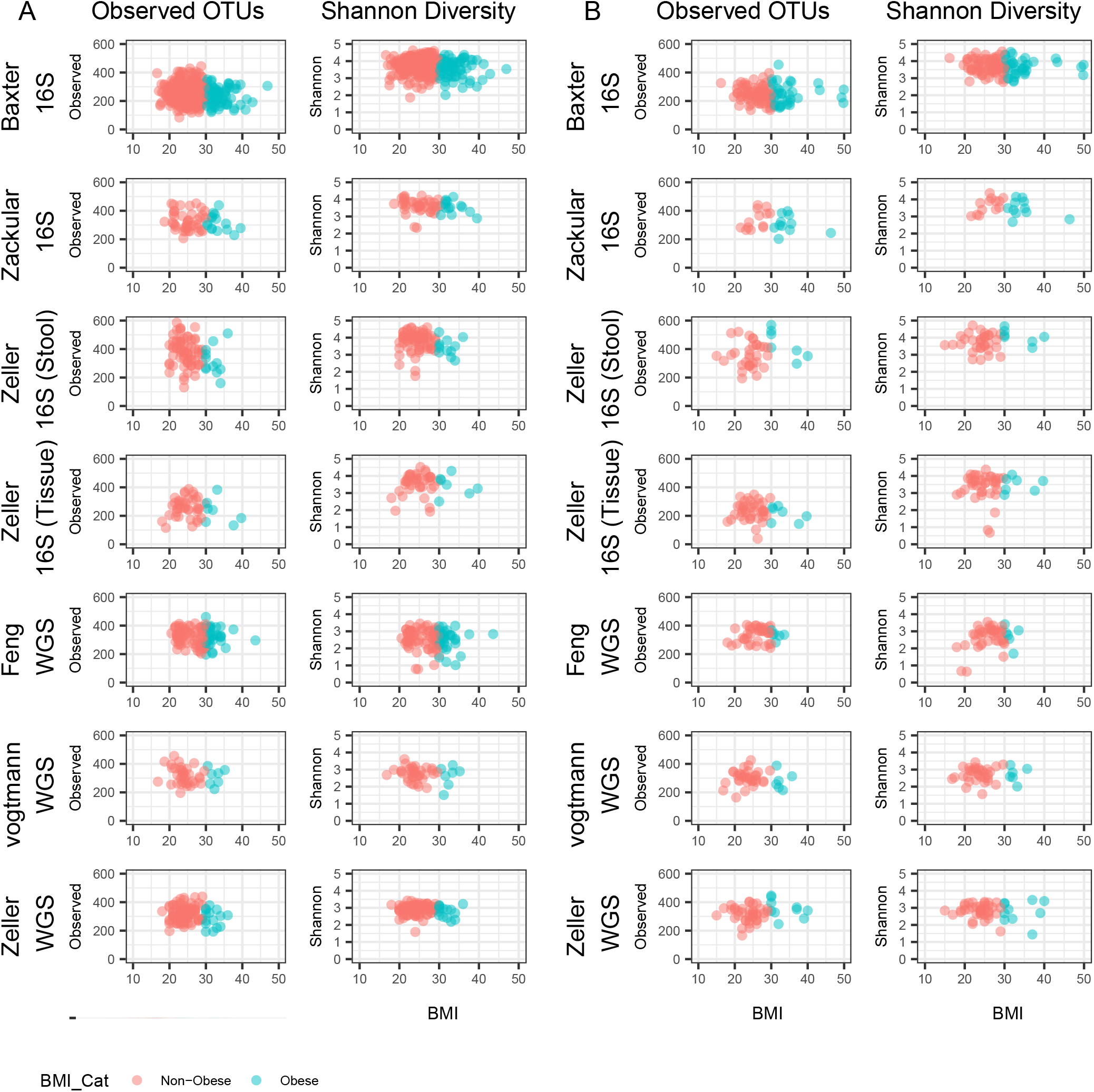
Alpha diversity by BMI in individuals with or without CRCA) Observed OTUs and Shannon diversity in individuals without CRC comparing BMI and alpha diversity metric, observed OTUs or Shannon diversity respectively. B)Observed OTUs and Shannon diversity in individuals with CRC comparing BMI and alpha diversity metric, observed OTUs or Shannon diversity respectively.

**Figure S3:**
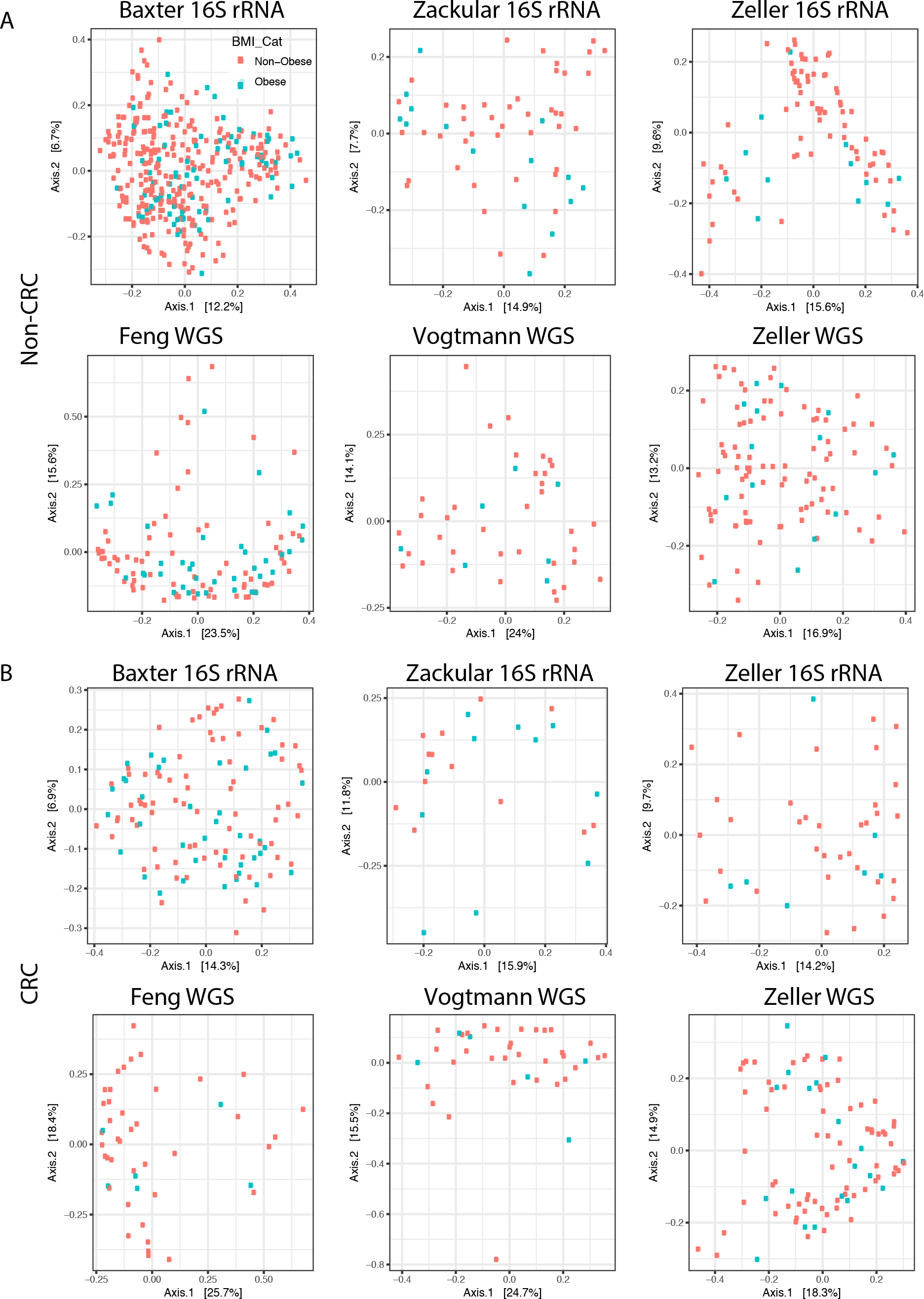
Beta diversity in induvial with or without obesity and with or without CRC A) Differences in community composition between individuals with and without obesity among those without CRC B) or with CRC. The axes were found using PCoA using Bray-Curtis distances among points with the proportion of variance accounted for by each axis reported. Points are colored by obesity status.

**Figure S4:**
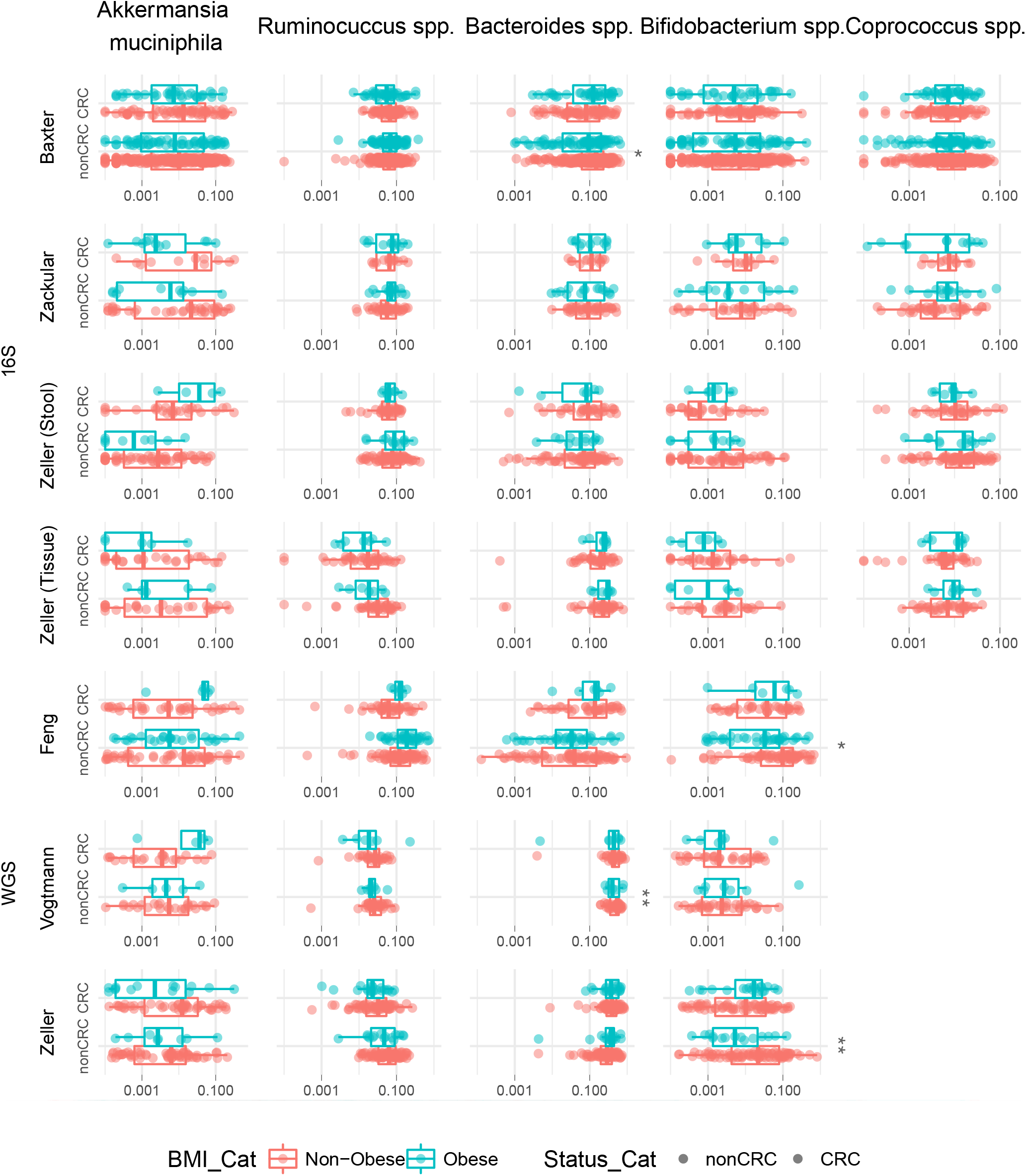
Differential abundance of taxa associated with obesity and CRC taxa.OTUs (16S rRNA) or species (WGS) log 10 scale relative abundance of *Ruminiococcus spp., Coprococcus spp., Bacteroides spp., Bifidobacterium spp*. and *Akkermansia muciniphila*. P-values were calculated using negative binomial regression using abundance as a count and including age and sex as covariates. Significant differences between obese v non-obese with or without CRC are denoted by an asterisk (FDR adjusted p-value <0.1)

**Figure S5:**
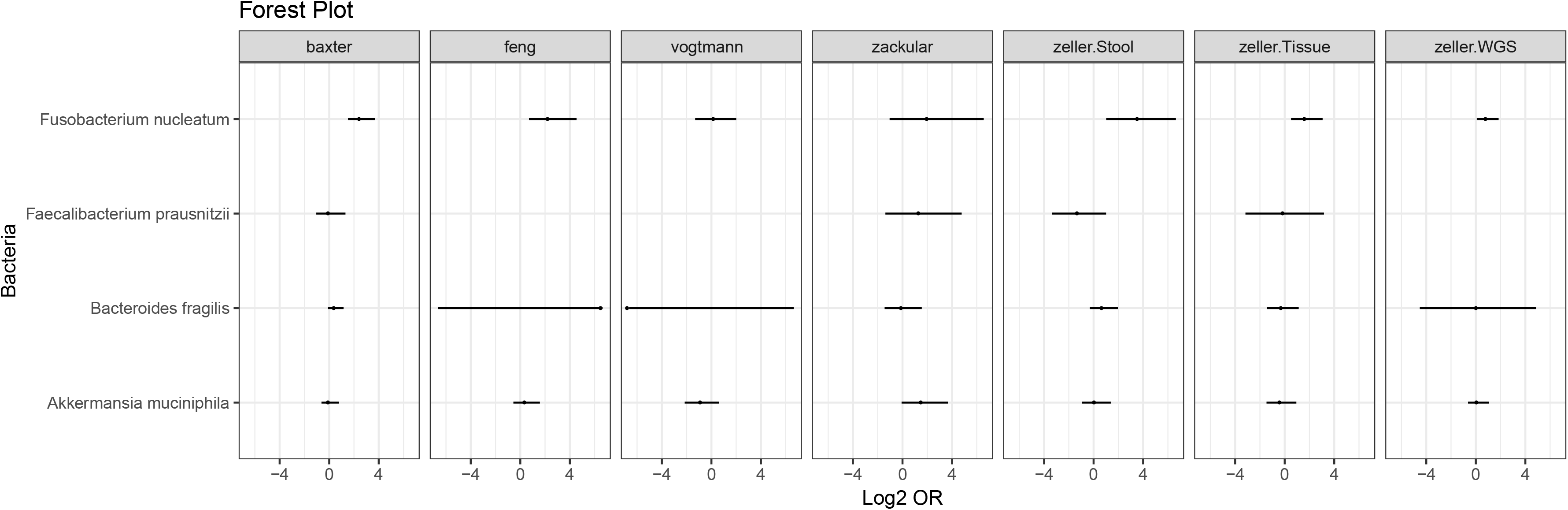
Ability of CRC-associated taxa to predict CRC among individuals with obesity. For each species identified from previous CRC microbiome studies, *F. nucleatum, F. prausnitzii, B. fragilis*, or *A. muciniphila*, the log_2_odd ratio was calculated for individuals with obesity to determine odds of being classified as having CRC.

**Supplemental Table 1:**
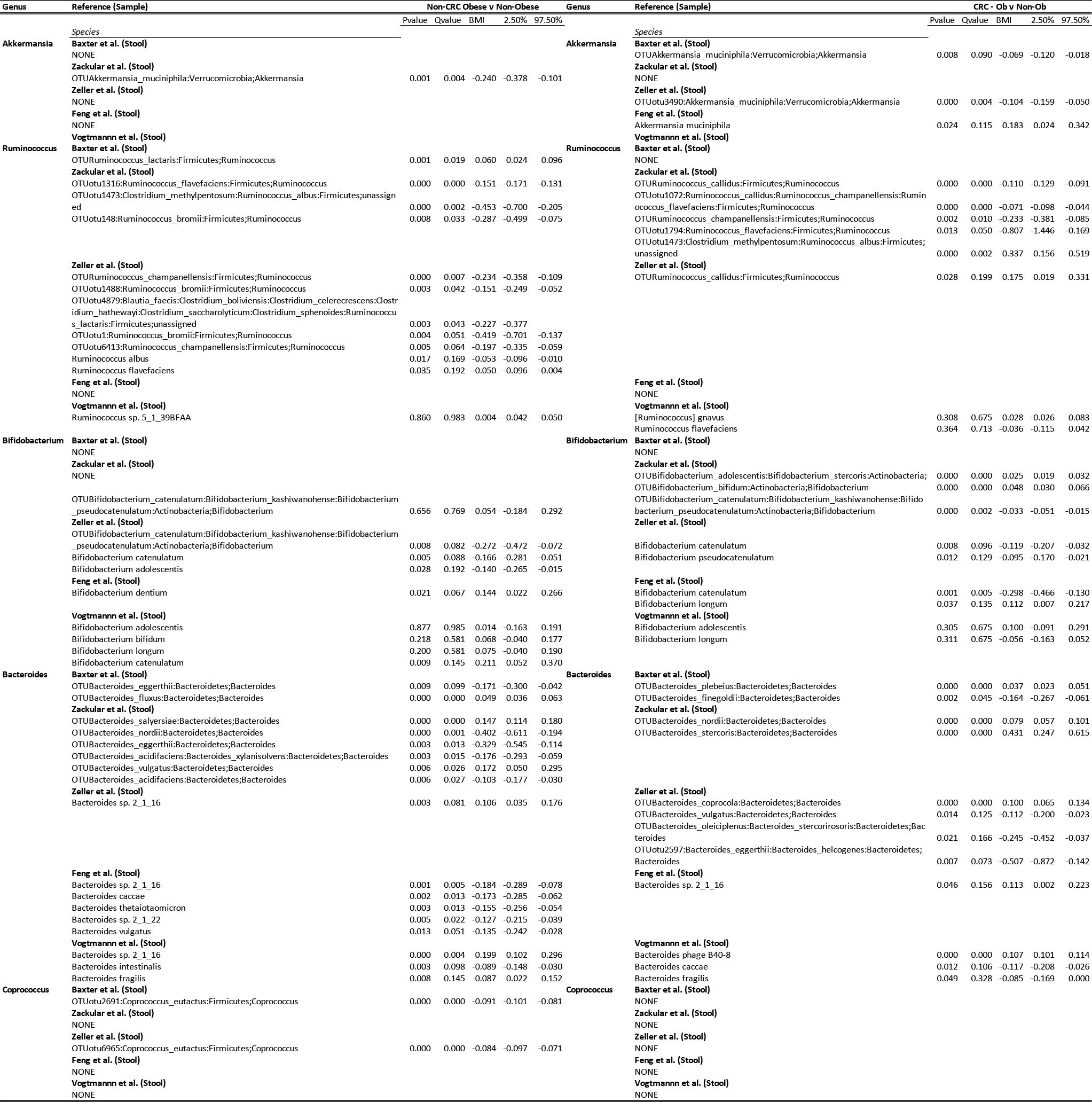
Summary of Species Level Differential Abundance.

**Supplemental Table 2:** Mediation analysis among taxa associatied with obesity and CRC.

| Study | Taxa | Change in |  |  |  |  |
| --- | --- | --- | --- | --- | --- | --- |
|  |  | Prob.<sup>a</sup> | Est | p-value | LL (2.5) | UL (97.5) |
| baxter | Bacteroides_fluxus | -0.00865 | 0.00562 | 0.14969 | -0.00203 | 0.01328 |
| baxter | Howardella_ureilytica | -0.00741 | 0.00067 | 0.60715 | -0.00188 | 0.00321 |
| baxter |  | -0.00722 |  |  |  |  |
|  | otu1526:Enterorhabdus_caecimuris:Enterorhabdus_mucosicola |  | -1.00E-04 | 0.85142 | -0.00117 | 0.00097 |
| baxter | otu2405:Allobaculum_stercoricanis | -0.00726 | 9.00E-05 | 0.87294 | -0.00106 | 0.00125 |
| baxter | otu2695:Eubacterium_coprostanoligenes | -0.00865 | 0.00562 | 0.30655 | -0.00515 | 0.01639 |
| baxter | otu2753:Vallitalea_guaymasensis | -0.00785 | 0.00243 | 0.64083 | -0.00779 | 0.01266 |
| baxter | otu847:Clostridium_aerotolerans:Clostridium_algidixylanolyticum:Clostridium_saccharolyticum:Clostridium_xylanolyticum:Gracilibacter_thermotolerans | -0.00798 | 0.00296 | 0.62175 | -0.00881 | 0.01474 |
| baxter | otu911:Intestinimonas_butyrificiproducens | -0.008 | 0.00304 | 0.66624 | -0.01077 | 0.01685 |
| feng | Haemophilus parainfluenzae | 0.00763 | -5.00E-05 | 0.98819 | -0.00621 | 0.00612 |
| feng | Lactobacillus casei group | 0.00753 | 0.00037 | 0.88939 | -0.00482 | 0.00556 |
| feng | Prevotella denticola | 0.00884 | -0.00487 | 0.7282 | -0.03235 | 0.0226 |
| feng | Prevotella ruminicola | 0.00878 | -0.00464 | 0.70382 | -0.02859 | 0.0193 |
| zackular | Bacteroides_eggerthii | -0.01117 | -0.00027 | 0.99158 | -0.05084 | 0.0503 |
| zackular | Bacteroides_nordii | -0.01306 | 0.0073 | 0.50709 | -0.01427 | 0.02887 |
| zackular | Bacteroides_salysiae | -0.01058 | -0.00262 | 0.62977 | -0.01325 | 0.00802 |
| zackular | Gemmiger_formicilis | -0.01153 | 0.00116 | 0.72148 | -0.0052 | 0.00751 |
| zackular | otu1157:Alistipes_indistinctus | -0.01127 | 0.00014 | 0.98768 | -0.01788 | 0.01816 |
| zackular | otu1202:Clostridium_botulinum:Clostridium_sporogenes | -0.01062 | -0.00248 | 0.69992 | -0.01512 | 0.01015 |
| zackular | otu1257:Intestinimonas_butyrificiproducens | -0.01126 | 9.00E-05 | 0.98154 | -0.00782 | 0.00801 |
| zackular | otu1316:Ruminococcus_flavefaciens | -0.01065 | -0.00236 | 0.74719 | -0.01668 | 0.01197 |
| zackular | otu1989:Eubacterium_coprostanoligenes | -0.01113 | -0.00041 | 0.90967 | -0.0075 | 0.00668 |
| zackular | otu2005:Alistipes_finegoldii:Alistipes_massiliensis | -0.01097 | -0.00108 | 0.75814 | -0.00793 | 0.00578 |
| zackular | otu327:Prevotella_oris | -0.01158 | 0.00136 | 0.77451 | -0.00794 | 0.01066 |
| zackular | otu476:Clostridium_saccharogumia | -0.01117 | -0.00026 | 0.96635 | -0.01256 | 0.01203 |
| zackular | otu508:Caloramator_fervidus:Trigonala_elaeagnus | -0.01108 | -0.00064 | 0.94102 | -0.01763 | 0.01635 |
| zackular | Phascolarctobacterium_succinatutens | -0.01117 | -0.00025 | 0.92066 | -0.00523 | 0.00473 |
| zeller.Stool | Clostridium_bolteae:Clostridium_clostridioforme | -0.00063 | 0.00049 | 0.87263 | -0.00547 | 0.00644 |
| zeller.Stool | Dialister_invisus | -0.00037 | -0.00052 | 0.94127 | -0.01446 | 0.01341 |
| zeller.Stool | Dialister_succinatophilus | -0.00081 | 0.00121 | 0.76375 | -0.0067 | 0.00912 |
| zeller.Stool | otu2321:Streptococcus_salivarius:Streptococcus_thermophilus:Streptococcus_vestibularis | 0.00046 | -0.00388 | 0.81746 | -0.03681 | 0.02906 |
| zeller.Stool | otu2483:Prevotella_copri | -9E-05 | -0.00167 | 0.80792 | -0.01516 | 0.01182 |
| zeller.Stool | otu4937:Peptococcus_niger | -0.00023 | -0.00108 | 0.76023 | -0.00805 | 0.00588 |
| zeller.Stool | otu834:Prevotella_copri:Prevotella_stercorea | -0.00049 | -5.00E-05 | 0.97294 | -0.00321 | 0.0031 |
| zeller.Stool | Phascolarctobacterium_succinatutens | -0.00054 | 0.00013 | 0.99755 | -0.08285 | 0.08311 |
| zeller.Stool | Streptococcus_porcinus:Streptococcus_seminale:Streptococcus_uberis | 0.00037 | -0.00352 | 0.62505 | -0.01762 | 0.01059 |
| zeller.Stool | Streptococcus_salivarius | 0.00138 | -0.00756 | 0.28432 | -0.02139 | 0.00628 |
| zeller.WGS | Bifidobacterium_bifidum | -0.00557 | 0.00212 | 0.98433 | -0.2097 | 0.21395 |
| zeller.WGS | Bifidobacterium_catulatum | -0.00443 | -0.00244 | 0.37338 | -0.00783 | 0.00294 |
| zeller.WGS | Streptococcus_salivarius | -0.00694 | 0.00761 | 0.41392 | -0.01065 | 0.02588 |
| zeller.WGS | Bacteroides sp. 2_1_16 | 0.05349 |  | 1.05494 |  |  |
| zeller.WGS | Streptococcus_salivarius | 0.05349 | 0.0257 | 1.05494 | 1.02604 | -0.0289 |
| zeller.WGS | [Eubacterium] eligens | 0.05349 |  | 1.05494 |  |  |
| zeller.WGS | Bifidobacterium_bifidum | 0.05349 | 0.0312 | 1.05494 | 1.03169 | -0.02325 |

<sup>a</sup> quantification of the change in OR from model 1 to model 2.

**Supplemental Table 3:** Mediation Cross Tabulation.

| Study | Bacteria | CRC_Bact_Present | CRC_Bact_NotPresent | nonCRC_Bact_Present | nonCRC_Bact_NotPresent |
| --- | --- | --- | --- | --- | --- |
| baxter | otu2695:Eubacterium_coprostanoligenes | 102 | 54 | 267 | 63 |
| baxter | otu911:Intestinimonas_butyrificiproducens | 209 | 87 | 160 | 30 |
| baxter | otu2405:Allobaculum_stercoricanis | 91 | 32 | 278 | 85 |
| baxter | Howardella_ureilytica | 70 | 28 | 299 | 89 |
| baxter | otu1526 | 83 | 28 | 286 | 89 |
| baxter | otu2753:Vallitalea_guaymasensis | 181 | 45 | 188 | 72 |
| baxter | otu847 | 237 | 60 | 132 | 57 |
| baxter | Bacteroides_fluxus | 65 | 3 | 304 | 114 |
| zackular | otu508:Caloramator_fervidus | 32 | 13 | 26 | 11 |
| zackular | Gemmiger_formicilis | 52 | 20 | 6 | 4 |
| zackular | otu1989:Eubacterium_coprostanoligenes | 21 | 9 | 37 | 15 |
| zackular | otu2005 | 9 | 4 | 49 | 20 |
| zackular | Phascolarctobacterium_succinatutens | 16 | 6 | 42 | 18 |
| zackular | otu1257:Intestinimonas_butyrificiproducens | 30 | 13 | 28 | 11 |
| zackular | Bacteroides_salysiae | 6 | 2 | 52 | 22 |
| zackular | otu476:Clostridium_saccharogumia | 44 | 17 | 14 | 7 |
| zackular | Bacteroides_eggerthii | 25 | 5 | 33 | 19 |
| zackular | otu1157:Alistipes_indistinctus | 25 | 9 | 33 | 15 |
| zackular | otu1202 | 10 | 4 | 48 | 20 |
| zackular | Bacteroides_nordii | 24 | 6 | 34 | 18 |
| zackular | otu327:Prevotella_oris | 15 | 8 | 43 | 16 |
| zackular | otu1316:Ruminococcus_flavifaciens | 13 | 8 | 45 | 16 |
| zeller.Stool | Phascolarctobacterium_succinatutens | 42 | 21 | 44 | 20 |
| zeller.Stool | otu4937:Peptococcus_niger | 18 | 11 | 68 | 30 |
| zeller.Stool | otu2483:Prevotella_copri | 29 | 13 | 57 | 28 |
| zeller.Stool | otu834 | 15 | 6 | 71 | 35 |
| zeller.Stool | Dialister_invisus | 29 | 10 | 57 | 31 |
| zeller.Stool | Streptococcus_salivarius | 9 | 1 | 77 | 40 |
| zeller.Stool | Streptococcus_porcinus | 6 | 2 | 80 | 39 |
| zeller.Stool | Dialister_succinatiphilus | 20 | 8 | 66 | 33 |
| zeller.Stool | Clostridium_bolteae | 53 | 26 | 33 | 15 |
| zeller.Stool | otu2321 | 58 | 21 | 28 | 20 |
| feng | Ruminococcus sp. 5_1_39BFAA | 110 | 46 |  |  |
| feng | Prevotella_denticola | 73 | 40 | 37 | 6 |
| feng | Lactobacillus_casei group | 81 | 36 | 29 | 10 |
| feng | Haemophilus_parainfluenzae | 84 | 36 | 26 | 10 |
| feng | Prevotella_ruminicola | 72 | 39 | 38 | 7 |
| feng | Coprobacillus sp. D7 | 110 | 46 |  |  |
| zeller.WGS | Bifidobacterium_catenumatum | 103 | 88 | 2 | 1 |
| zeller.WGS | Bacteroides sp. 2_1_16 | 105 | 89 |  |  |
| zeller.WGS | Streptococcus_salivarius | 105 | 88 |  |  |
| zeller.WGS | [Eubacterium] eligens | 105 | 89 |  |  |
| zeller.WGS | Bifidobacterium_bifidum | 105 | 88 |  |  |

ium\_aerotolerans:Clostridium\_algidixylanolyticum:Clostridium\_saccharolyticum:Clostridium
s\_thermophilus:Streptococcus\_vestibularis otu508:Caloramator\_fervidus:Trigonala\_elaeagnus Clostridium\_bolteae:Clostridium\_clostridioforme Streptococcus\_porcinus:Streptococcus\_seminale:Streptococcus\_uberis lla\_copri:Prevotella\_stercorea

**Supplemental Table 4:** Taxa with Q values <0.2 after performing mediation analysis between PooledDS and PooledBMI.

| **Baxter** | **Zackular** | **Zeller (16S)** | **Feng** | **Zeller (WGS)** | **Vogtmann** |
| --- | --- | --- | --- | --- | --- |
| OTUotu2695 | OTUotu508 | OTUPhascolarctobacterium_succinatutens | Ruminococcus sp. 5_1_39BFAA | Bifidobacterium catenulatum | None |
| OTUotu911 | OTUGemmiger_formicilis | OTUotu4937 | Prevotella denticola | Bacteroides sp. 2_1_16 |  |
| OTUotu2405 | OTUotu1989 | OTUotu2483 | Lactobacillus casei group | Streptococcus salivarius |  |
| OTUHowardella_ureilytica | OTUotu2005 | OTUotu834 | Haemophilus parainfluenzae | [Eubacterium] eligens |  |
| OTUotu1526 | OTUPhascolarctobacterium_succinatutens | OTUDialister_invisus | Prevotella ruminicola | Bifidobacterium bifidum |  |
| OTUotu2753 | OTUotu1257 | OTUStreptococcus_salivarius | Coprobacillus sp. D7 |  |  |
| OTUotu847 | OTUBacteroides_salyersiae | OTUStreptococcus_porcinus |  |  |  |
| OTUBacteroides_fluxus | OTUotu476 | OTUDialister_succinatiphilus |  |  |  |
|  | OTUBacteroides_eggerthii | OTUClostridium_bolteae |  |  |  |
|  | OTUotu1157 | OTUotu2321 |  |  |  |
|  | OTUotu1202 |  |  |  |  |
|  | OTUBacteroides_nordii |  |  |  |  |
|  | OTUotu327 |  |  |  |  |
|  | OTUotu1316 |  |  |  |  |

